# Dynamic High-Content Imaging Reveals Surface Exposure of Virulent *Leishmania* Amastigotes in Infected Macrophages Undergoing Pyroptosis

**DOI:** 10.1101/2020.02.03.931907

**Authors:** Thibault Rosazza, Hervé Lecoeur, Thierry Blisnick, Maryse Moya-Nilges, Pascale Pescher, Phillipe Bastin, Eric Prina, Gerald F. Späth

**Affiliations:** Institut Pasteur, Unité de Parasitologie Moléculaire et Signalisation, INSERM U1201, Paris, France; Institut Pasteur International Mixed Unit ‘Inflammation and *Leishmania* infection’; Institut Pasteur, Trypanosome Cell Biology Unit & INSERM U1201, Paris, France; Institut Pasteur, Unité de Technologie et service BioImagerie Ultrastructurale (UtechSPBI), INSERM 1201, Paris, France

**Keywords:** *Leishmania*, macrophage, pyroptosis, high-content, single-cell, real-time imaging, virulence

## Abstract

*Leishmania* spp are obligate intracellular parasites that infect vertebrate phagocytes, notably macrophages. We previously reported that *Leishmania amazonensis* (*L. am*) subvert the host cell pro-inflammatory response by dampening the macrophage NLRP3 inflammasome. No information is available on how *Leishmania* infection affects inflammatory cell death termed pyroptosis, known to limit microbial infection. Here, we provide first evidence that *L. amazonensis*-infected macrophages can undergo pyroptosis when subjected to pro-inflammatory stimuli. We analyzed the dynamics of the pyroptotic process and the fate of intracellular amastigotes at the single cell level using spinning disk confocal microscopy and high-content, real-time imaging. Bone marrow-derived macrophages (BMDMs) were infected with *L. am* amastigotes isolated from footpad lesions and sequentially treated with lipopolysaccharide (LPS) and adenosine triphosphate (ATP) for canonical NLRP3 inflammasome priming and activation. Real-time monitoring was performed for 240 min post ATP addition. Longitudinal analyses revealed distinct phases of the pyroptotic process, including rapid decay of the parasitophorous vacuole (PV) as monitored by the pH-sensitive lysotracker fluid phase marker, progressive decrease in macrophage viability as monitored by accumulation of the nuclear dye YO-PRO-1, followed by translocation of the luminal PV membrane to the cell surface observed for 40% of macrophages, resulting in the extracellular exposure of amastigotes that remained anchored to the PV membrane. Scanning and transmission electron microscopy analyses revealed a highly polarized orientation of parasites with exclusive exposure of the anterior pole toward the extracellular milieu, and an attachment site forming a potential biological junction between the parasite posterior pole and the PV membrane. We showed that the exposed parasites are resistant to the cytolytic host cell activities linked to pyroptosis and retain their full infectious potential in reinfection experiments using naïve macrophages. Together these data uncover a novel *Leishmania* immune subversion strategy that may allow stealthy parasite dissemination via the uptake of pyroptotic host debris by uninfected phagocytes.

## Introduction

*L. amazonensis (L. am)* is one of the etiologic agents of human, localized cutaneous and anergic diffuse cutaneous leishmaniasis in South America (Barral et al., 1991; Silveira et al., 2004). Restriction of intracellular *L. am* replication *in vitro* and *in vivo* was shown to depend on the NLRP3 (NOD-, LRR-and pyrin domain-containing protein 3) inflammasome (Zamboni and Sacks, 2019). This intracellular sensor is induced in response to a “priming signal” represented by cytokines or ligands of Toll-like receptors, and further activated by damage-associated molecular patterns (DAMPs), such as ATP. Inflammasome activation triggers caspase-1 activity, which cleaves pro-IL-1β and pro-IL-18 into mature cytokines - promoting an anti-microbial, pro-inflammatory immune response (Swanson et al., 2019) - and cleaves gasdermin D that forms pores inside the macrophage plasma membrane, sustaining IL-1β release (Lieberman et al., 2019) and causing pyroptosis (Bergsbaken et al., 2009; Fink and Cookson, 2006; Jorgensen et al., 2017; Shi et al., 2015). Significantly, pyroptotic cell death contributes to a protective response against intracellular pathogens by removing the niche for infection (Bergsbaken et al., 2009), exposing microbes to the immune system (Jorgensen and Miao, 2015), rendering them more susceptible to anti-microbial agents (Jorgensen et al., 2016) or directly killing them via the gasdermin D lytic activity (Liu et al., 2016).

Even though the interaction of *L. am* with the NLRP3 inflammasome has attracted considerable attention (Lecoeur, 2019; Lima-Junior et al., 2013), how pyroptosis affects parasite survival and virulence remains to be elucidated. Here, we deployed a real-time, high-content single-cell analysis that uncover unique features of *Leishmania* amastigotes during host macrophage pyroptosis.

## Materials & Methods

### Ethics statement

Animals were housed at the Institut Pasteur animal facilities accredited by the French Ministry of Agriculture for performing experiments on live rodents. Work on animals was performed in compliance with French and European regulations on care and protection of laboratory animals (EC Directive 2010/63, French Law 2013-118, February 6th, 2013). All experiments were approved by the Ethic Committee for animal experimentation (CETEA#89) and authorized by the French ministry of higher education, research and innovation under the reference 2013-0092 in accordance with the Ethics Charter of animal experimentation that includes respect of the 3Rs principles, appropriate procedures to minimize pain and animal suffering.

### Bone Marrow-Derived Macrophage cultures

Female C57BL/6 mice were obtained from Janvier (Saint Germain-sur-l’Arbresle, France). Bone marrow cell suspensions were recovered from tibias and femurs as described (Courret et al., 1999). Bone marrow cells were plated at 1.5×10^7^ cells/mL in hydrophobic Petri dishes (Corning Life Science – 664161) and cultured for 6 days at 37°C in a 7.5% CO_2_ air atmosphere in complete DMEM medium (Pan Biotech – P04-03500) containing 4.5 g/L Glucose, 2mM L-Glutamine, 1mM sodium pyruvate and 3.7 g/L NaHCO3 and supplemented with 15% FCS (Gibco, A3160801), 10 mM HEPES (Gibco, 15630080), 50-μg/mL Penicillin/Streptomycin (Sigma, P4333), 50 μM 2-Mercaptoethanol (Sigma, M6250) and with 75 ng/mL recombinant mouse CSF-1 (Immunotools, 1234311). Adherent bone marrow-derived macrophages (BMDMs) were recovered after a treatment with 25 mM EDTA (Sigma) for 30 min at 37°C. BMDMs were seeded in complete DMEM medium supplemented with 30 ng/mL rmCSF-1 into various culture-treated supports, including (i) flat bottom 96-well black plates (Sigma, 655090) for OPERA analyses, (ii) 24-well plates (Falcon, 353047) with glass coverslips inside for scanning electron microscopy analyses, and (iii) 6-well plates (Falcon, 353046) for amastigote isolation from infected BMDMs.

### Macrophage infection and activation

*Leishmania amazonensis* amastigotes (LV79 strain, WHO - MPRO/BR/72/M1841) expressing mCherry (Lecoeur, 2019) were isolated from footpad lesions of infected Swiss nu/nu mice and purified as described (Courret et al., 1999). Infections were carried out at 34°C at a ratio of four amastigotes per macrophage. *Leishmania donovani* amastigotes (strain 1S2D, MHOM/SD/62/1S-CL2D) were isolated from infected spleens of female RjHan:AURA Golden Syrian hamsters and purified as described (Pescher et al., 2011; Prieto Barja et al., 2017). Infections were carried out at 37°C at a ratio of eight amastigotes per macrophage.

All pyroptosis experiments were performed after 3 days of infection using a sequential treatment of 500 ng/ml LPS (Alpha Diagnostic, LPS11-1) for 4 hrs and 5 mM ATP (Sigma, A26209) for different periods up to 4 hours.

### Real time analysis of pyroptosis by confocal microscopy

The following fluorescent reporters were added to the cell cultures 15 mn before ATP stimulation: Hoechst 33342 (10 μg/mL) (Invitrogen, H3570), LysoTracker Green DND-26 (1 μM) (Invitrogen, L7526), YO-PRO-1 (1 μM) (Invitrogen, Y3603). The pyroptotic process was monitored at 34°C and 7.5% CO_2_ using a fully automated spinning disk confocal microscope (PerkinElmer Technologies, OPERA QEHS) with a 40x water immersion objective (Aulner et al., 2013). Image acquisition was performed every five minutes after ATP addition using the following sequential acquisition settings: (i) 405 nm laser line excitation, filter 450/50 for Hoechst 33342 detection, (ii) 488 nm laser line excitation, filter 540/75 for Lysotracker DND-26 or YO-PRO-1 detection and (iii) 561 nm laser line excitation, filter 600/40 for *mCherry* detection. Additionally, transmission light microscopy images were taken for analyses of the macrophage PV area and amastigote localization. Fifteen fields at the same focal plane were taken every 5 minutes for every channels and for each sample. Images were transferred to the Columbus Conductor^Tm^ Database (Perkin Elmer Technologies) for storage and further analysis using specific scripts (Fig S1A and B) or exported for analysis using the FIJI software.

### Epifluorescence microscopy analysis of pyroptotic macrophages

Pyroptotic BMDMs were fixed for 1 hour at room temperature (RT) in 4% paraformaldehyde (PFA) (Electron Microscopy Sciences, 15710), then incubated with PBS containing 50 mM NH_4_Cl (Sigma, A9434) for 10 min and washed in the same buffer. Samples were incubated in PBS containing 10 μg/mL of anti-mouse CD68 (Biolegend, 137001) or IgG2a isotype control (Biolegend, 400501) and 0.25% of gelatin for 1 hour at RT. After a washing step, samples were incubated in a solution containing 10 μg/mL of anti-Rat FITC conjugated antibody (Jackson ImmunoResearch, 712-096-153) and 0.25% of gelatin for 1 hour at RT. Finally, samples were washed in PBS, incubated in PBS containing 10 μg/mL of Hoechst 33342 (Invitrogen, H3570) for 10 min at RT, and coverslips were mounted in Mowiol following further washing steps in PBS and distilled water. Images were acquired using the UltraView VOX microscope and data were analysed using the FIJI software.

### Scanning and transmission electron microscopy analyses

Pyroptotic BMDMs were fixed overnight with 2.5% glutaraldehyde in 0.1 M cacodylate buffer (pH 7.2) at 4°C and post-fixed in 0.1 M cacodylate buffer (pH 7.2) containing 1% OsO4. After serial dehydration, samples were critical-point dried (Emitech K850 or Balzers Union CPD30) and coated with gold using a sputter coater (Gatan Ion Beam Coater 681). Scanning electron microscopy observations were made with the JEOL 7600F microscope. Images were colorized using Adobe Photoshop CS software.

For transmitted electron microscopy analysis, cells were cultured on coverslips and were fixed with 2.5 % of glutaraldehyde (Sigma Aldrich) in PHEM buffer (120 mM PIPES, 50 mM HEPES, 20 mM EGTA, 4mM MgCl2, pH 7.3). Post fixation was done with 1 % osmium tetroxide (EMS 19152 2% aqueous solution, MERK) and 1.5 % ferrocyanide (Sigma Aldrich 8131) in PHEM. After dehydration by a graded series of ethanol from 25 to 95% grade 1, the samples were infiltrated with epoxy resin as described (Loussert et al., 2012). The 70 nm sections obtained by thin sectioning using a Leica UC 7 microtome (Leica microsystems), were collected on formvar coated slot grids (EMS 215-412-8400) and were contrasted with 4% uranyl acetate and Reynolds lead citrate (Delta microscopies, 11300). Stained sections were observed with a Tecnai spirit FEI operated at 120 kV. Images were acquired with FEI Eagle digital camera.

### Amastigote isolation from pyroptotic and control macrophages

Supernatants of BMDM cultures (pyroptotic and control BMDMs) were carefully replaced by fresh medium without LPS/ATP. After scraping of adherent macrophages using a plastic cell scrapper (Greiner bio-one, 541070) and multiple passages through a 27 gauge needle to free amastigotes from cell remnants, isolated amastigotes were centrifuged, washed in PBS, counted and used to assess parasite viability, the capacity of amastigotes to differentiate into proliferating promastigotes *in vitro*, and amastigote virulence by infection of naïve BMDMs.

### Flow cytometric analysis of parasite death by YO-PRO-1 staining

Amastigotes isolated from Leishmania-infected, pyroptotic and control BMDMs were seeded in a 96-well plate (Falcon, 353072) at a final concentration of 5×10^6^ parasites per mL and incubated for 10 min with YO-PRO-1^™^ iodide at 0.1 μM final concentration (Thermo Fisher Scientitic, Y3603). Samples were then immediately analysed on the CytoFLEX cytometer (Beckman Coulter) in a BSL2 containment to evaluate YO-PRO-1 incorporation. Data were analysed using the Kaluza software package. Lesion-derived amastigotes treated with 70% EtOH for 10 minutes were included as positive control for cell death and YO-PRO-1 incorporation.

### Analysis of parasite differentiation, growth and metabolic status

Amastigote to promastigote differentiation was analyzed by SEM (as described above) directly on amastigotes exposed on pyroptotic macrophages after 24 hours incubation in 1 mL of promastigote culture medium at 27°C. For growth analysis in culture, purified amastigotes were incubated at 27°C in promastigote culture medium at 10^5^ parasites / mL. Promastigote culture density was determined daily during 6 consecutive days by visual counting using a Malassez chamber. The parasite metabolic status was analyzed at days 2 and 3 by resazurin assay. Briefly, 200 μl of parasite culture was transferred into wells of a 96-well plate (Sigma, 655090) and incubated with 2.5 μg/mL of resazurin (Sigma, R7017) for 4 hours at 27°C. The fluorescence intensity of the resazurin-derived resorufin was determined using the Tecan Safire2 plate reader (558 nm excitation, 585 nm emission) (Lamotte et al., 2019).

### Determination of amastigote virulence

Amastigote virulence was assessed using two protocols. First, isolated amastigotes from *Leishmania*-infected, pyroptotic and control BMDMs were added at a ratio of four parasite per macrophages on naive BMDMs seeded on glass coverslips placed in 24-well plates. One and two days after incubation at 34°C, amastigote-loaded BMDMs were fixed with 4% PFA (Electron Microscopy Sciences, 15710) and incubated for 10 min in PBS containing 10 μg/mL of Hoechst 33342 (Thermo Fisher Scientitic, H3570). Image acquisition was performed with the Axio Imager (Carl Zeiss) (405 nm laser line excitation, filter 450/50). The number of parasites per macrophage was visually determined by counting, considering at least one hundred macrophages. Second, the culture medium of pyroptotic macrophages was carefully replaced by pre-warmed BMDM medium containing freshly differentiated, naïve BMDMs (1:1 ratio between naïve and pyroptotic BMDMs). Three days later, live-images combining bright field and mCherry fluorescence were acquired using the EVOS microscope (Thermo Fisher Scientific) and analyzed with the FIJI software for infection of the naïve BMDMs.

## RESULTS AND DISCUSSION

### A high-content imaging protocol for dynamic single-cell analysis of pyroptotic macrophages infected by *L. amazonensis* amastigotes

To analyze the dynamics of pyroptosis in *L. am*- infected macrophages, we established a new imaging protocol combining high-content and single-cell analyses using the OPERA QEHS confocal plate reader (Fig. 1A). Bone marrow-derived macrophages (BMDMs) were seeded into 96 well plates (macrophage settlement). They were subsequently infected for three days with mCherry transgenic, virulent *L. am* amastigotes isolated from lesions of infected Nude mice, allowing to form characteristic communal PVs (parasite settlement). Then, LPS was added for 225 min (NLRP3 priming) followed by staining with fluorescent reporters. Pyroptosis was triggered after NLRP3 activation by ATP (pyroptosis induction). Dynamic cellular changes were monitored at high-content and single-cell levels by using Hoechst 33342 (for cell number, Fig. 1B1), YO-PRO-1 (for loss of plasma membrane (PM) integrity, Fig. 1B2) (Adamczak et al., 2014), LysoTracker green (LTG) (for PV acidity, Fig. 1B3) (Aulner et al., 2013), and mCherry (for parasite localization, figure 1B4) (Aulner et al., 2013). Macrophage morphology was analyzed by transmission light microscopy (TL, Fig. 1B5). High-content analyses (HCA) were carried out in real-time up to 240 min post ATP addition. Analyses at the population level were performed using segmentation procedures of the Columbus^™^ Image Data Storage and Analysis system (Figs S1A, S1B). The analysis of fluorescent read-outs at 240 min documented that (i) no macrophages were lost during the analyses (Fig. 1C1), (ii) pyroptosis induction was successful (Fig. 1C2), and the PV integrity was lost in all pyroptotic cells (Figure 1C3). Single-cell analyses using the ImageJ software package permitted to determine macrophage / PV area and parasite location (Fig. 1D).

**FIGURE 1:**
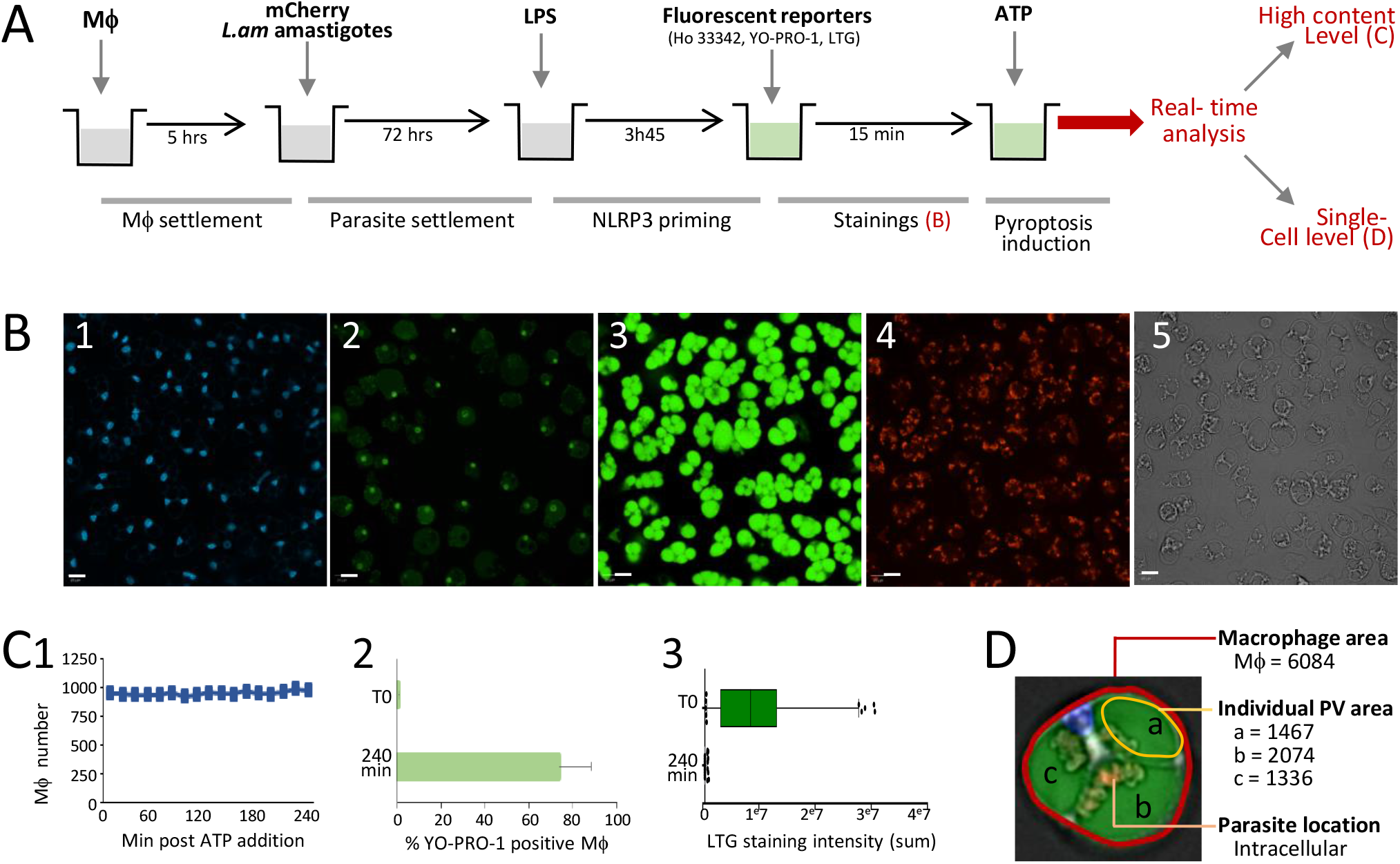
Design of a real-time imaging assay to monitor amastigote location, PV features and pyroptosis in *L. amazonensis*-infected BMDMs. **A: Experimental flow chart** Mouse BMDMs were derived from bone marrow progenitors in presence of mCSF1 and seeded in black flat bottom, tissue culture-treated 96-well plates. Five hours later, mCherry lesion-derived amastigotes were added to BMDMs (MOI = 4:1, 72 hrs, 34°C). NLRP3 was primed adding 500 ng/ml LPS for 3h 45 min. Then fluorescent reporters Hoechst 33342 (Ho 33342), LysoTracker Green DND-26 (LTG) and YO-PRO-1 were added. Finally, pyroptosis was triggered by NLRP3 activation with 5 mM ATP. Real-time analyses were performed using the OPERA QEHS confocal plate reader at 34°C, 7.5% CO_2_ for 240 min. **B: High Content Analysis read-outs** Image acquisition was performed every 5 minutes. A representative field image is displayed for each channel (T0 time point for Ho 33342, LTG, mCherry and Transmitted Light (TL); T = 120 min for YO-PRO-1 staining). Macrophage nuclei were stained by Hoechst 33342 (blue channel, B1). Nuclei of pyroptotic macrophages were stained by YO-PRO-1 (green channel, B2). PV integrity and acidity were evaluated by the LTG staining (green channel, B3). Parasite location was determined by the mCherry fluorescence signal (red channel, B4). Macrophage morphology was evaluated by cell area (TL, B5). Scale bar = 20 μm. **C: HCA quality controls** HCA analyses were performed at the population level analyzing 15,000 cells per well. Three quality controls were performed by: i) monitoring macrophage numbers to control that no cells were lost during the analysis (C1), ii) analyzing the percentage of dead macrophages (YO-PRO-1 positive) 4 hours post ATP addition to control for efficient pyroptosis induction (C2, at least 70% of death must be observed at this time point), and iii) assessing LTG fluorescence at T0 and T240 to monitor PV presence (T0) and pyroptosis associated with the complete loss of PV staining (T240, C3). **D: Single cell analyses** Merged pictures (Ho 33342, LTG, mCherry and TL images) of representative infected macrophages (T0). Identification and quantification of areas of PVs (a), (b) and (c) and parasite location were performed using the Image J software. The unit of PV area is square pixels (s.p.).

### Rapid decay of PV precedes amastigote extracellular exposure during macrophage pyroptosis

Pyroptosis dynamics, PV integrity and parasite localization were analyzed every 5 min after ATP addition for a duration of 240 min in infected, LPS-primed macrophages. HCA of YO-PRO-1 incorporation revealed that pyroptosis was asynchronous as judged by the progressive increase in dead cells (Fig. 2A, black curve). Pyroptosis occurred with a constant rate (+18.2 ±4.2% dead cells per hour). A loss of LTG staining revealed that PVs rapidly decayed (Fig. 2A, green curve), following three distinct empirical rates: (i) a fast decay rate during the first 45 minutes (−4.2e6 ± 1.2e6 / 30 min, stage 1) leading to a 47.6% decrease of LTG fluorescence intensity, (ii) an intermediate rate (45 to 120 minutes, −1.2e6 ± 6.3e5 / 30 min, stage 2), and (iii) a slow rate thereafter (0.58e6 ± 0.37e6 / 30 min, stage 3). This decay may result from ATP-induced changes of the vacuolar pH, and of PV integrity that could be caused by osmotic changes similar to those described for lysosomes (Guha et al., 2013; Takenouchi et al., 2009). Single-cell analyses revealed differences in the PV decay kinetics between macrophages (Fig. S2) as well as between PVs inside a same macrophage (Fig. 2B1 - 3).

**FIGURE 2:**
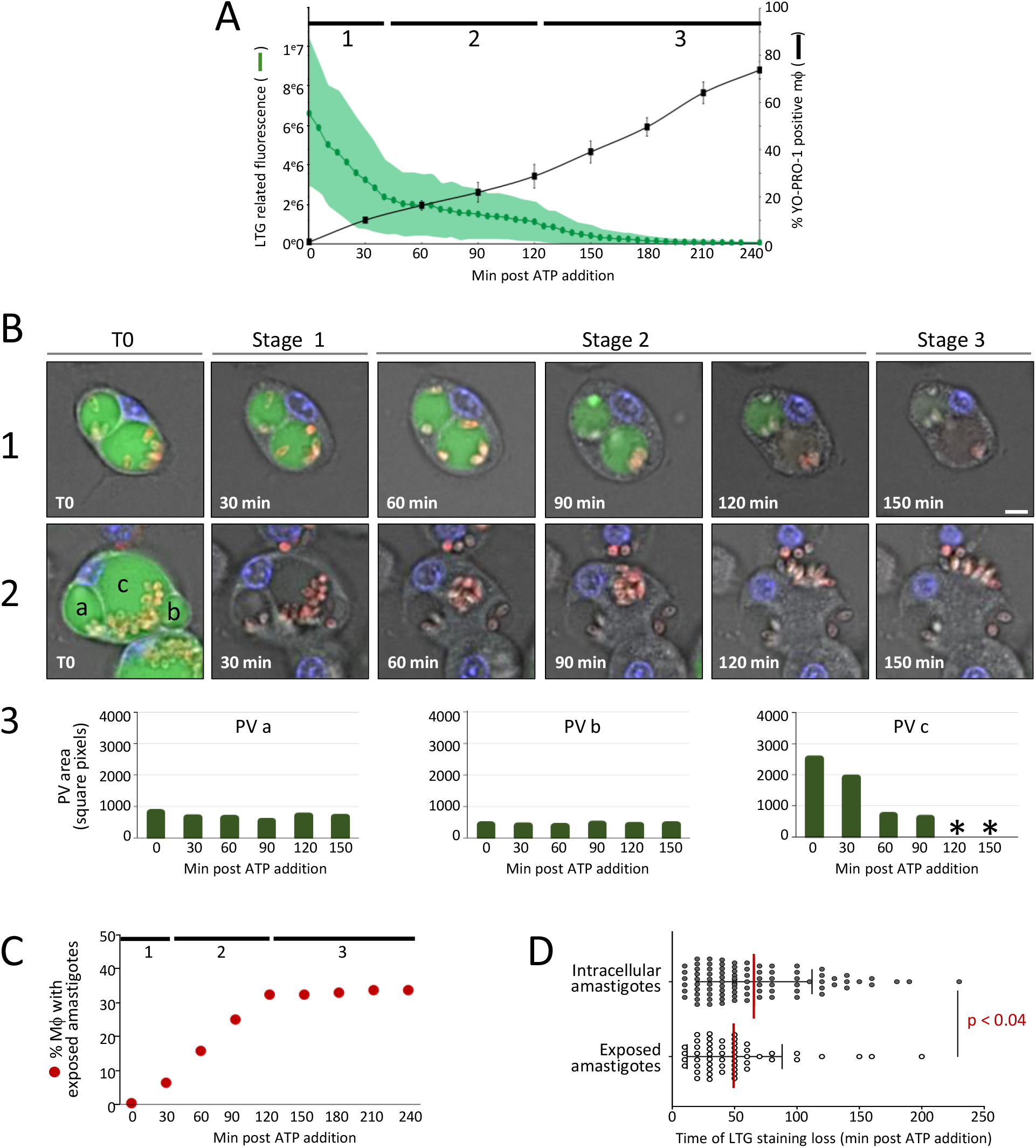
Real-time multiparametric analysis of pyroptotic, *L. amazonensis*-infected macrophages. Pyroptosis was induced in *L. amazonensis-infected* BMDMs by LPS and ATP. Real-time analyzes were performed using the QEHS High Content Imaging System to follow macrophage pyroptosis features and amastigote location during 240 minutes. **(A) High content analysis of PV acidity and macrophage pyroptosis.** Macrophage pyroptosis (mean YO-PRO-1 positive macrophages ± SEM, black curve) and PV acidity (mean LTG fluorescence ± SEM, green curve and green area, respectively, n= 2 independent experiments) were monitored. Three stages were delineated according to the decay rate of LTG fluorescence. **(B) Real-time, single cell monitoring of infected macrophages during pyroptosis.** Merged images for Hoechst 33342, LTG, mCherry fluorescence and TL acquired from two representative macrophages during the first 150 min. Acquisition was performed before ATP addition (T0) and during the pyroptosis stages defined in figure 2A1. (B1) Macrophage displaying 2 PVs with different LTG decay rates and maintaining amastigotes intracellularly. (B2) Macrophage displaying 3 PVs (a, b, c), with one exposing amastigotes to the extracellular milieu (c). (B3) Area of PVs (a), (b) and (c) from the macrophage displayed in B2. Scale bar = 20 μm. **(C) Single cell monitoring of surface amastigote exposition during pyroptosis.** The percentage of macrophages exposing amastigotes at their surface was determined every 30 min (single cell analysis, n= 384 macrophages). The three pyroptosis stages are indicated. **(D) Relationship between amastigote location and LTG staining decay.** The time point at which LTG staining is lost is indicated for macrophages with intracellularly trapped (black dots) or surface exposed (white dots) amastigotes. The statistical significance between the 2 groups is indicated (p value).

HCA analysis revealed that amastigotes were not released into the culture medium (Fig. S2): They either remained intravacuolar (62% of macrophages) or were exposed at the cell surface (38% of macrophages) as shown in representative cells (Fig. 2B1 and 2). Significantly, different decay rates and parasite localizations were observed for individual PVs, even inside the same macrophage as indicated in figure 2B2, showing three independent vacuoles (a, b and c) that rapidly lost LTG fluorescence (stage 1). PVs (a) and (b) retained their size and their amastigotes intracellularly (Fig. 2B2, 3) whereas PV (c) collapsed (60 min) and the luminal side of its membrane was exposed to the extracellular milieu (120 min, stage 2). PV lumen exposure could be triggered by ATP as judged by previous reports implicating ATP in the extracellular discharge of secretory exosomes (Andrei et al., 2004; Bergsbaken et al., 2011), in exosome exocytosis (Qu et al., 2007), in the release of autophagolysosomes and phagocytosed, intracellular particles (Bergsbaken et al., 2011; Takenouchi et al., 2009). Amastigote exposure was confirmed by Scanning Electronic Microscopy (Fig. S3) and occurred preferentially in macrophages displaying a rapid PV decay (Fig. 2B, C, D). Parasite exposure did not correlate with PV size, PV number nor amastigote number per vacuole (data not shown).

### Ultrastructural analysis of amastigote attachment zone and surface exposure

Pyroptosis has been recognized as an anti-microbial strategy during bacterial infection, capable of pathogen trapping inside dying host cells (Jorgensen et al., 2016) and directly killing bacteria by the lytic activity of Gasdermin D (Liu et al., 2016). Our observation that amastigotes remain attached on the surface of pyroptotic cells primed us to investigate parasite integrity and the molecular structures underlying parasite surface retention.

Scanning Electron Microscopy analyses (SEM) showed that 38.8 % of pyroptotic macrophages displayed intact parasites at the cell surface (Fig. S3A), thus confirming our results obtained with the OPERA system (Fig. 2C). Ultrastructural analysis demonstrated that exposed amastigotes were trapped in blebbing macrophage membrane structures that are similar to previously described membrane shedding associated with IL-1β secretion (Pizzirani et al., 2007). Significantly, exposed amastigotes showed a highly polarized orientation with only the anterior pole (flagellar side) being exposed toward the extracellular milieu (Fig. 3A1, 2). This feature was also observed for pyroptotic macrophages infected with *L. donovani* amastigotes, that reside in individual PVs (Fig. S3C). We next investigated the posterior attachment zone by Transmission Electron Microscopy (TEM), which revealed an electro-dense, membranous junction formed between parasite and PV membranes both in non-stimulated and pyroptotic macrophages (Fig. 3B1, 3B2). This junction corresponds to a defined attachment site - showing features similar to gap junctions - that permits amastigotes of communal *Leishmania* species to be anchored to PV membranes (Benchimol and de Souza, 1981). We next performed epifluorescence microscopy analysis of CD68, a transmembrane molecule located specifically in lysosomal compartments (Chistiakov et al., 2017), including PVs (Antoine et al., 1998). During pyroptosis, amastigotes remained embedded in CD68 positive PV membranes, either on the surface (Figure 3C2, yellow arrow) or inside dead macrophages (Fig. 3C2, white arrow).

**FIGURE 3:**
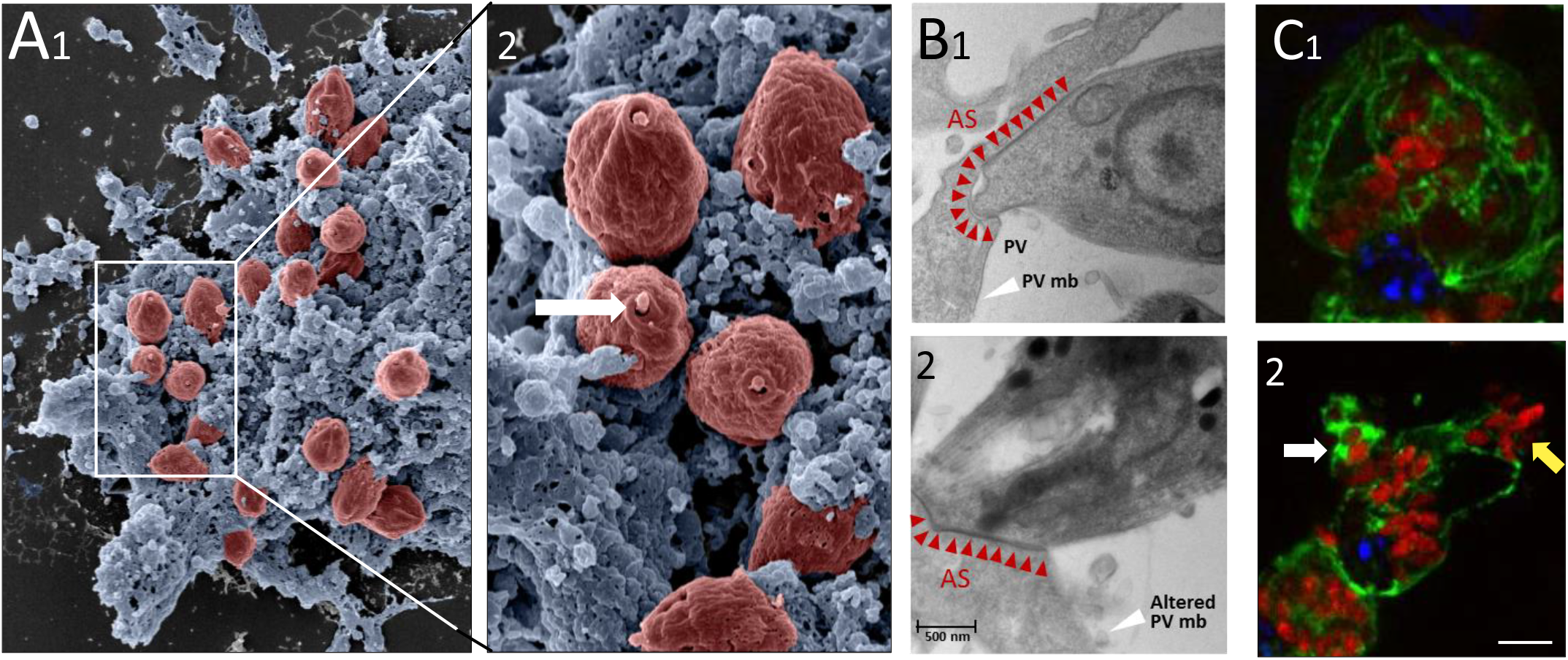
Amastigote exposure during macrophage pyroptosis. Pyroptosis was induced in *L.am*-infected BMDMs by LPS and ATP stimulation. (A) SEM analysis of a representative pyroptotic macrophage exposing amastigotes (red colorization) at the cell surface (1). Zoomed picture (2) highlights the unique orientation of attached parasites with the flagellar pocket pointing towards the extracellular milieu (white arrow). (B) TEM images showing the Attachment Site (AS, red arrowheads) of an amastigote in untreated (1) and pyroptotic (2) macrophages. Note the strong unaltered PV membrane at the AS in the pyroptotic macrophage that contrasts with altered nearby membranes. White arrowheads point to the PV membrane. “PV” indicates the PV lumen. (C) Epifluorescence microscopy analysis of CD68 staining (green channel), mCherry-parasites (red channel), and Hoechst 33342-stained macrophage nucleus (blue channel). BMDMs were analyzed before (1) and after ATP addition (2). White and yellow arrows indicate amastigotes embedded in compact CD68^+^ membranes or attached to CD68^+^ membranes but exposed to the extracellular milieu, respectively.

These results demonstrate that amastigotes remain attached to pyroptotic macrophages through strong interactions between parasite and host membranes at their attachment site (see also Movie 1), which represents a new type of pathogen trapping in pyroptotic cells that differs from those previously described (Jorgensen et al., 2016).

### *Leishmania amazonensis* amastigotes are resistant to host cell pyroptosis and retain full infectivity

Since bacteria can be damaged by gasdermin D during host cell pyroptosis (Liu et al., 2016), we next investigated whether parasite viability and infectivity were affected during macrophage pyroptosis. Amastigotes isolated from pyroptotic macrophages (A-pm) were compared to amastigotes isolated from unstimulated control macrophages (A-cm). Macrophage pyroptosis did not reduced parasite viability (99% of A-pm remained YO-PRO-1 negative, Fig. 4A1). Second, when pyroptotic macrophages were cultured in promastigote-specific medium (48 hours, 26°C), amastigotes rapidly transformed into ovoid flagellated, motile and proliferating promastigotes that detached from macrophage remnants (Fig. 4A2). These promastigotes showed normal growth (visual parasite counting, Fig. 4A3) and metabolic activity (resazurin reduction data, Fig. 4A4). Finally, A-pm maintained their capacity to establish efficient new infections (Fig. 4B). In addition, amastigotes associated to macrophage remnants were efficiently phagocytosed by naïve BMDMs and established normal infection levels as shown by the formation of typical large communal PVs housing numerous parasites after a 3 day co-culture (Fig. 4C3).

**FIGURE 4:**
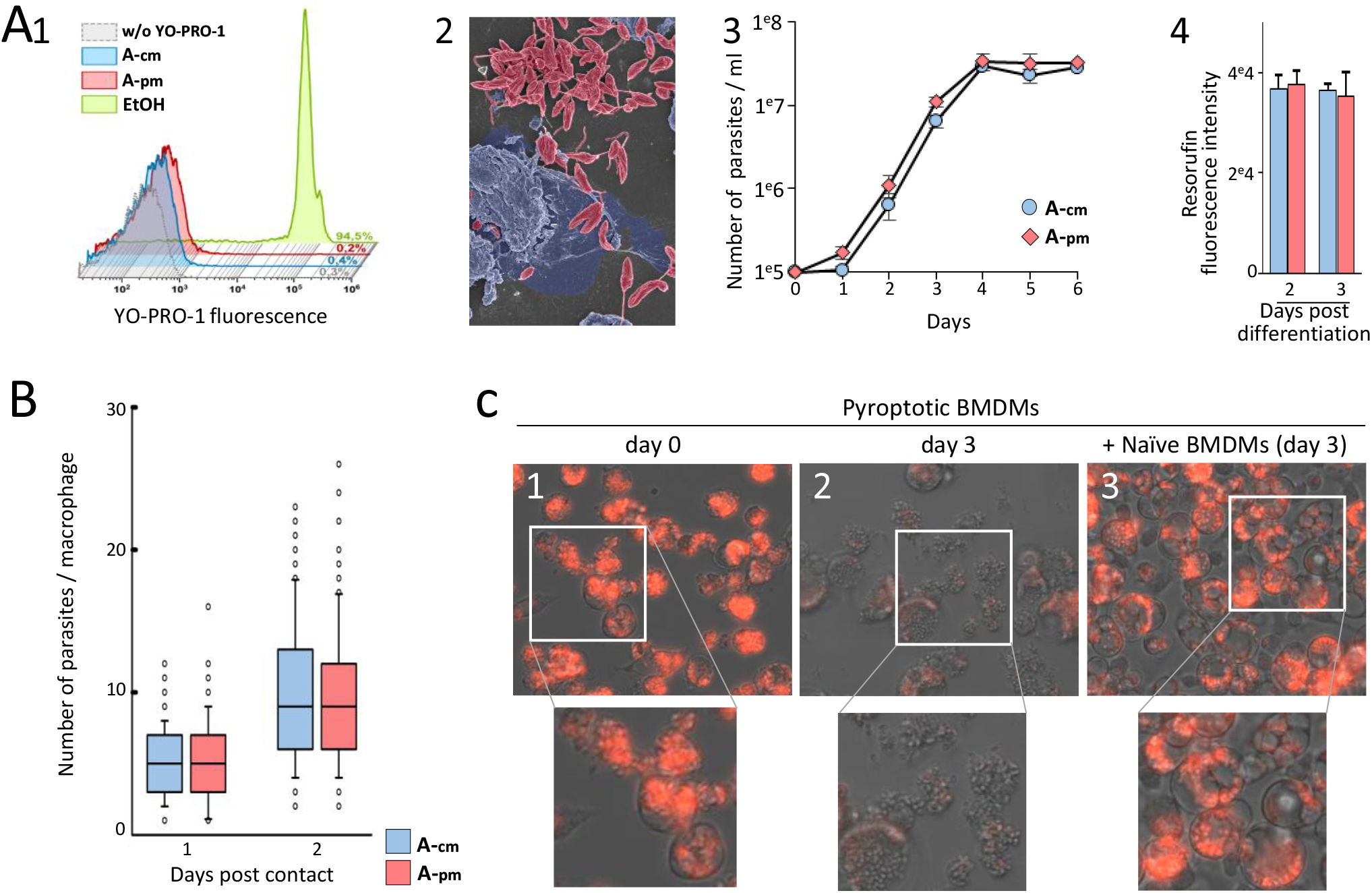
Amastigotes derived from pyroptotic macrophages are viable and virulent. Parasites isolated from pyroptotic (A-pm) and control unstimulated (A-cm) macrophages were compared. (A) Assessment of viability and amastigote-to-promastigote differentiation capacity. (1) FACS analysis of YO-PRO-1 incorporation of isolated amastigotes and ethanol-killed parasites (positive control for cell death). Percentages of YO-PRO-1^+^ parasites are indicated. (2) SEM pseudo-colored image of a representative pyroptotic macrophage 4 hours after pyroptosis induction and maintained in promastigote medium (27°C, 48 hours). Promastigotes were colored in red. (3) Growth curves for A-pm and A-cm (27°C). (4) Assessment of metabolic activity of A-pm-and A-cm-derived promastigotes (raw resorufin fluorescence values are shown). (B) Evaluation of amastigote virulence. A-pm and A-cm were added to cultures of naïve BMDMs for one and two days. Parasite number per macrophage is shown (n = 2 independent experiments). (C) Evaluation of the capacity of amastigotes attached to pyroptotic remnants to infect naïve macrophages. Naïve BMDMs were added to pyroptotic *L*. am-infected macrophages in fresh medium (34°C, 3 days) and analysed by epifluorescence and phase contrast microscopy. Representative fields and a cropped zoomed area are shown (upper and lower panels, respectively). Pyroptotic macrophages at day 0 (1) and 3 (2). The loss of mCherry fluorescence was probably due to parasite death in absence of host cells. (3) Pyroptotic macrophages at day 3 post addition of naïve macrophages (Representative pictures, n = 2 independent experiments).

## CONCLUSION

Here the dynamic interactions between *Leishmania* amastigotes and PVs in pyroptotic BMDMs were analyzed at the population-, single-cell and ultrastructural levels. Our real-time HCA uncovered three distinct pyroptosis phases: During stage 1, PV membrane decays (rapid loss of PV acidity and reduced PV size). In stage 2, the PV lumen is externalized in 38% of cells, inducing exposure of infectious, membrane-anchored amastigotes (Fig. S4). In stage 3, cellular alterations are more sustained (80% cell death). *L. am* amastigotes displayed the unique ability to preserve their viability and infectivity during macrophage pyroptosis. Consequently, *in vivo*, parasite trapping to pyroptotic debris may favor parasite spreading - as shown for other forms of host cell death – allowing for stealthy phagocytosis by freshly recruited macrophages in the inflammatory infection site (de Menezes et al., 2016). On the other hand, the extracellular exposure of amastigotes could expose parasites to complement-dependent cytotoxicity that is known to control parasite load *in vivo* (Laurenti et al., 2004), or to anti-leishmanial antibodies that could promote cell-mediated cytotoxicity via Fc receptor-dependent phagocytosis. Our results will stimulate future studies designed to assess the role of macrophage pyroptosis in *Leishmania* dissemination, transmission, and immune-pathology.

## Legend to supplementary figures

**SUPPLEMENTARY FIGURE 1: Automatic image analysis procedure to monitor macrophage pyroptosis and PV features in *L. amazonensis-infected* BMDMs.**

Live imaging was performed with the OPERA plate reader every 5 minutes following pyroptosis induction (ATP addition). Images were transferred to the Columbus Conductor^™^ Database (Perkin Elmer Technologies) and analyzed using the integrated Image analysis building blocks. The sequential image segmentation process showing the identification and counting of Hoechst-stained living/dead macrophages are shown for YO-PRO-1-stained dying cells (A) and LTG-stained PVs (B).

**SUPPLEMENTARY FIGURE 2: Characteristics of PVs and amastigotes during pyroptosis**. A representative field of infected macrophages showing 3 time points corresponding to the 3 stages defined in figure 2A was followed during the pyroptotic process. The merged fluorescence and bright field images illustrate the contrasting behavior of LTG-stained PVs and mCherry-expressing amastigotes in Hoechst 33342-stained macrophages. Note the different location of amastigotes, either within PV (black arrow) or exposed to the outside milieu (white arrow) at the 240 min time point.

**SUPPLEMENTARY FIGURE 3: Exposed amastigotes are attached through their posterior pole to pyroptotic macrophages**

Scanning Electron Microscopy (SEM) was performed on *L. amazonensis* (A, B) and *L. donovani* (C)-infected BMDMs. (A) Representative image of a macrophage population during pyroptosis after 3 hours of ATP stimulation. Amastigotes are attached to dead macrophages and exposed to the extracellular milieu (red colorization). (B) Representative image of an untreated, living and healthy macrophage population. Note the presence of numerous big globular cells corresponding to heavily infected macrophages harboring large PVs. (C) Cropped and zoomed image showing a representative pyroptotic *L. donovani*-infected BMDM. Note that the exposition at the cell surface of *L. donovani* amastigotes (olive colorization) is similar to that of *L. amazonensis* parasites and reveal the parasite flagellar pocket pointing towards the extracellular milieu (arrow).

**SUPPLEMENTARY FIGURE 4: Synthetic graphical abstract recapitulating the fate of intracellular *Leishmania* during macrophage pyroptosis**.

The triggering of pyroptosis in *Leishmania*-infected macrophages by ATP leads to two different scenarios. In both cases, PVs decay with a loss of LTG fluorescence, with or without shrinkage: In 61% of macrophages, a slow reduction of LTG staining occurs, with PV maintained *in cellula* and intracellular trapping of virulent amastigotes (upper part). In 39% of macrophages, PVs decay faster and the inner part of the PV membrane is exposed on the macrophage surface (Exposure of the Luminal side of the parasitophorous Vacuole, ELV). ELV leads to the polarized exposure of virulent amastigotes that are maintained embedded in PV-membranes (lower part). Parasite exposure occurs only during the first two stages of pyroptosis. The third stage is associated to more pronounced cellular alterations.

## Acknowledgements

We thank Drs Jacomina Krijnse Locker and Geneviève Milon for scientific discussions and Dr Nathalie Aulner for help with the Opera system.

## Funding

This project was supported by i) a fund of the Institut Pasteur International Direction (International Mixed Unit ‘Inflammation and Leishmania infection’), ii) the French National Research Agency (ANR-10-INSB-04-01, Investments for the Future), the Conseil de la Region Ile-de-France (program Sesame 2007, project Imagopole, S. Shorte) and the Fondation Française pour la Recherche Médicale (Programme Grands Equipements) (UtechS PBI/C2RT), and iii) a French Government Investissement d’Avenir programme, Laboratoire d’Excellence “Integrative Biology of Emerging Infectious Diseases” (ANR-10-LABX-62-IBEID) (Trypanosome Cell Biology Unit).

## Competing interests

Authors have no conflict of interest.

## Author contributions

TR, MMN, TB and PP performed experiments and analysed the data. TR, HL and EP designed the experiments, and analysed the data. HL, EP and GFS wrote the manuscript. PB planned EM analyses, scientific discussions, and correction of the manuscript.

## Bibliography

Adamczak, S.E., de Rivero Vaccar¡, J.P., Dale, G., Brand, F.J., 3rd, Nonner, D., Bullock, M.R., Dahl, G.P., Dietrich, W.D., and Keane, R.W. (2014). Pyroptotic neuronal cell death mediated by the AIM2 inflammasome. Journal of cerebral blood flow and metabolism: official journal of the International Society of Cerebral Blood Flow and Metabolism 34, 621–629.

Andrei, C., Margiocco, P., Poggi, A., Lotti, L.V., Torrisi, M.R., and Rubartelli, A. (2004). Phospholipases C and A2 control lysosome-mediated IL-1 beta secretion: Implications for inflammatory processes. Proceedings of the National Academy of Sciences of the United States of America 101, 9745–9750.

Antoine, J.C., Prina, E., Lang, T., and Courret, N. (1998). The biogenesis and properties of the parasitophorous vacuoles that harbour Leishmania in murine macrophages. Trends in microbiology 6, 392–401.

Aulner, N., Danckaert, A., Rouault-Hardoin, E., Desrivot, J., Helynck, O., Commere, P.H., Munier-Lehmann, H., Spath, G.F., Shorte, S.L., Milon, G., et al. (2013). High content analysis of primary macrophages hosting proliferating Leishmania amastigotes: application to anti-leishmanial drug discovery. PLoS neglected tropical diseases 7, e2154.

Barral, A., Pedral-Sampaio, D., Grimaldi Junior, G., Momen, H., McMahon-Pratt, D., Ribeiro de Jesus, A., Almeida, R., Badaro, R., Barral-Netto, M., Carvalho, E.M., et al. (1991). Leishmaniasis in Bahia, Brazil: evidence that Leishmania amazonensis produces a wide spectrum of clinical disease. The American journal of tropical medicine and hygiene 44, 536–546.

Benchimol, M., and de Souza, W. (1981). Leishmania mexicana amazonensis: attachment to the membrane of the phagocytic vacuole of macrophages in vivo. Zeitschrift fur Parasitenkunde 66, 25–29.

Bergsbaken, T., Fink, S.L., and Cookson, B.T. (2009). Pyroptosis: host cell death and inflammation. Nature reviews Microbiology 7, 99–109.

Bergsbaken, T., Fink, S.L., den Hartigh, A.B., Loomis, W.P., and Cookson, B.T. (2011). Coordinated host responses during pyroptosis: caspase-1-dependent lysosome exocytosis and inflammatory cytokine maturation. Journal of immunology 187, 2748–2754.

Chistiakov, D.A., Killingsworth, M.C., Myasoedova, V.A., Orekhov, A.N., and Bobryshev, Y.V. (2017). CD68/macrosialin: not just a histochemical marker. Laboratory investigation; a journal of technical methods and pathology 97, 4–13.

Courret, N., Prina, E., Mougneau, E., Saraiva, E.M., Sacks, D.L., Glaichenhaus, N., and Antoine, J.C. (1999). Presentation of the Leishmania antigen LACK by infected macrophages is dependent upon the virulence of the phagocytosed parasites. European journal of immunology 29, 762–773.

de Menezes, J.P., Saraiva, E.M., and da Rocha-Azevedo, B. (2016). The site of the bite: Leishmania interaction with macrophages, neutrophils and the extracellular matrix in the dermis. Parasites & vectors 9, 264.

Fink, S.L., and Cookson, B.T. (2006). Caspase-1-dependent pore formation during pyroptosis leads to osmotic lysis of infected host macrophages. Cellular microbiology 8, 1812–1825.

Jorgensen, I., and Miao, E.A. (2015). Pyroptotic cell death defends against intracellular pathogens. Immunological reviews 265, 130–142.

Jorgensen, I., Rayamajhi, M., and Miao, E.A. (2017). Programmed cell death as a defence against infection. Nature reviews Immunology 17, 151–164.

Jorgensen, I., Zhang, Y., Krantz, B.A., and Miao, E.A. (2016). Pyroptosis triggers pore-induced intracellular traps (PITs) that capture bacteria and lead to their clearance by efferocytosis. The Journal of experimental medicine 213, 2113–2128.

Lamotte, S., Aulner, N., Spath, G.F., and Prina, E. (2019). Discovery of novel hit compounds with broad activity against visceral and cutaneous Leishmania species by comparative phenotypic screening. Scientific reports 9, 438.

Laurenti, M.D., Orn, A., Sinhorini, I.L., and Corbett, C.E. (2004). The role of complement in the early phase of Leishmania (Leishmania) amazonensis infection in BALB/c mice. Brazilian journal of medical and biological research = Revista brasileira de pesquisas medicas e biologicas 37, 427–434.

Lecoeur, H. (2019). Leishmania targets the macrophage epigenome and dampens the NF-ĸB/NLRP3-mediated inflammatory response. bioRxiv.

Lieberman, J., Wu, H., and Kagan, J.C. (2019). Gasdermin D activity in inflammation and host defense. Science immunology 4.

Lima-Junior, D.S., Costa, D.L., Carregaro, V., Cunha, L.D., Silva, A.L., Mineo, T.W., Gutierrez, F.R., Bellio, M., Bortoluci, K.R., Flavell, R.A., et al. (2013).Inflammasome-derived IL-1beta production induces nitric oxide-mediated resistance to Leishmania. Nature medicine 19, 909–915.

Liu, X., Zhang, Z., Ruan, J., Pan, Y., Magupalli, V.G., Wu, H., and Lieberman, J. (2016). Inflammasome-activated gasdermin D causes pyroptosis by forming membrane pores. Nature 535, 153–158.

Loussert, C., Forestier, C.L., and Humbel, B.M. (2012). Correlative light and electron microscopy in parasite research. Methods in cell biology 111, 59–73.

Pescher, P., Blisnick, T., Bastin, P., and Spath, G.F. (2011). Quantitative proteome profiling informs on phenotypic traits that adapt Leishmania donovani for axenic and intracellular proliferation. Cellular microbiology 13, 978–991.

Pizzirani, C., Ferrari, D., Chiozzi, P., Adinolfi, E., Sandona, D., Savaglio, E., and Di Virgilio, F. (2007). Stimulation of P2 receptors causes release o IL-1beta-loaded microvesicles from human dendritic cells. Blood 109, 3856–3864.

Prieto Barja, P., Pescher, P., Bussotti, G., Dumetz, F., Imamura, H., Kedra, D., Domagalska, M., Chaumeau, V., Himmelbauer, H., Pages, M., et al. (2017). Haplotype selection as an adaptive mechanism in the protozoan pathogen Leishmania donovani. Nature ecology & evolution 1, 1961–1969.

Qu, Y., Franchi, L., Nunez, G., and Dubyak, G.R. (2007). Nonclassical IL-1 beta secretion stimulated by P2X7 receptors is dependent on inflammasome activation and correlated with exosome release in murine macrophages. Journal of immunology 179, 1913–1925.

Shi, J., Zhao, Y., Wang, K., Shi, X., Wang, Y., Huang, H., Zhuang, Y., Cai, T., Wang, F., and Shao, F. (2015). Cleavage of GSDMD by inflammatory caspases determines pyroptotic cell death. Nature 526, 660–665.

Silveira, F.T., Lainson, R., and Corbett, C.E. (2004). Clinical and immunopathological spectrum of American cutaneous leishmaniasis with special reference to the disease in Amazonian Brazil: a review. Memorias do Instituto Oswaldo Cruz 99, 239–251.

Swanson, K.V., Deng, M., and Ting, J.P. (2019). The NLRP3 inflammasome: molecular activation and regulation to therapeutics. Nature reviews Immunology 19, 477–489.

Takenouchi, T., Nakai, M., Iwamaru, Y., Sugama, S., Tsukimoto, M., Fujita, M., Wei, J., Sekigawa, A., Sato, M., Kojima, S., et al. (2009). The activation of P2X7 receptor impairs lysosomal functions and stimulates the release of autophagolysosomes in microglial cells. Journal of immunology 182, 2051–2062.

Zamboni, D.S., and Sacks, D.L. (2019). Inflammasomes and Leishmania: in good times or bad, in sickness or in health. Current opinion in microbiology 52, 70–76.

