## Supplemental figure 1A for "Dynamic High-Content Imaging Reveals Surface Exposure of Virulent *Leishmania* Amastigotes in Infected Macrophages Undergoing Pyroptosis"

### Slide 1
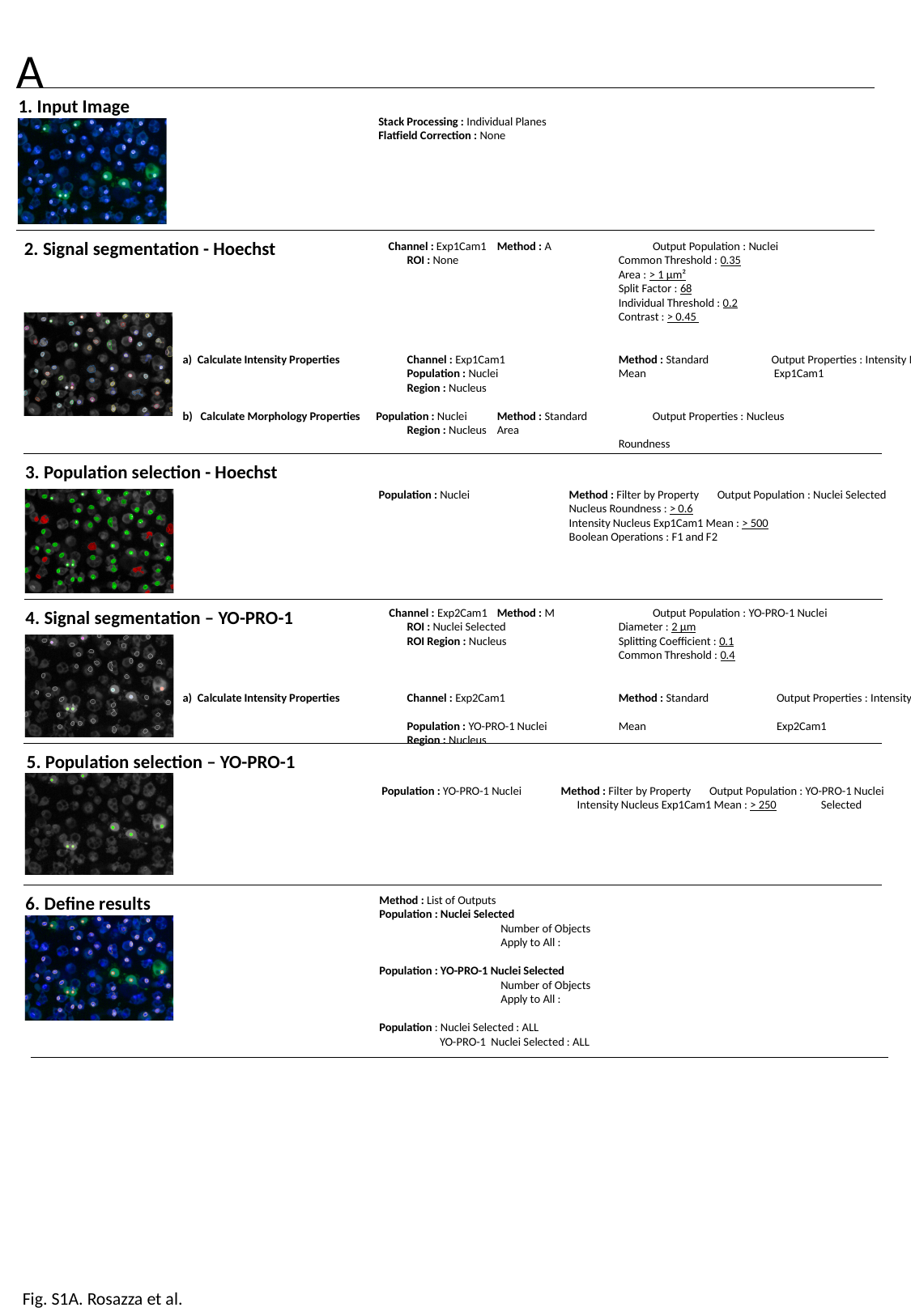

A
1. Input Image
Stack Processing : Individual Planes
Flatfield Correction : None
2. Signal segmentation - Hoechst
	 Channel : Exp1Cam1 	Method : A	 Output Population : Nuclei
		 ROI : None 		Common Threshold : 0.35
				Area : > 1 µm²
				Split Factor : 68
				Individual Threshold : 0.2
				Contrast : > 0.45
 a) Calculate Intensity Properties	 Channel : Exp1Cam1 	Method : Standard 	 Output Properties : Intensity Nucleus
		 Population : Nuclei	Mean	 Exp1Cam1
		 Region : Nucleus
 b) Calculate Morphology Properties Population : Nuclei 	Method : Standard 	 Output Properties : Nucleus
		 Region : Nucleus 	Area
				Roundness
3. Population selection - Hoechst
 Population : Nuclei 	 Method : Filter by Property Output Population : Nuclei Selected
	 Nucleus Roundness : > 0.6
	 Intensity Nucleus Exp1Cam1 Mean : > 500
	 Boolean Operations : F1 and F2
	 - Channel : Exp2Cam1 	Method : M	 Output Population : YO-PRO-1 Nuclei
		 ROI : Nuclei Selected 	Diameter : 2 µm
		 ROI Region : Nucleus	Splitting Coefficient : 0.1
				Common Threshold : 0.4
 a) Calculate Intensity Properties	 Channel : Exp2Cam1 	Method : Standard 	 Output Properties : Intensity Nucleus
		 Population : YO-PRO-1 Nuclei	Mean	 Exp2Cam1
		 Region : Nucleus
4. Signal segmentation – YO-PRO-1
5. Population selection – YO-PRO-1
 Population : YO-PRO-1 Nuclei Method : Filter by Property Output Population : YO-PRO-1 Nuclei
	 Intensity Nucleus Exp1Cam1 Mean : > 250 Selected
6. Define results
Method : List of Outputs
Population : Nuclei Selected
	Number of Objects
	Apply to All :
Population : YO-PRO-1 Nuclei Selected
	Number of Objects
	Apply to All :
Population : Nuclei Selected : ALL
 YO-PRO-1 Nuclei Selected : ALL
Fig. S1A. Rosazza et al.
