## Supplemental figure 1B for "Dynamic High-Content Imaging Reveals Surface Exposure of Virulent *Leishmania* Amastigotes in Infected Macrophages Undergoing Pyroptosis"

### Slide 1
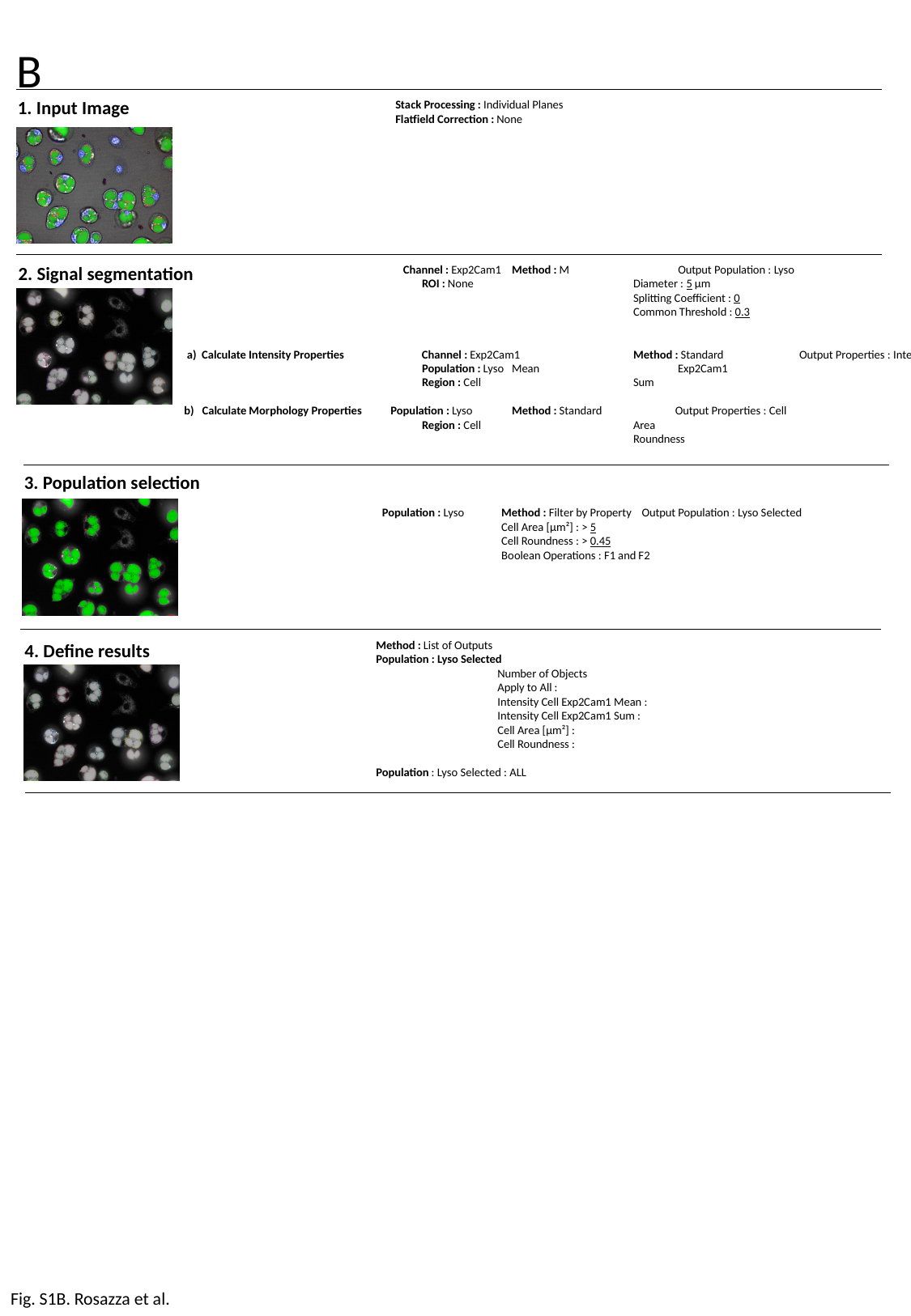

B
1. Input Image
Stack Processing : Individual Planes
Flatfield Correction : None
2. Signal segmentation
	 Channel : Exp2Cam1 	Method : M	 Output Population : Lyso
		 ROI : None 		Diameter : 5 µm
				Splitting Coefficient : 0
				Common Threshold : 0.3
 a) Calculate Intensity Properties	 Channel : Exp2Cam1 	Method : Standard 	 Output Properties : Intensity Cell
		 Population : Lyso	Mean	 Exp2Cam1
		 Region : Cell		Sum
 b) Calculate Morphology Properties Population : Lyso 	Method : Standard 	 Output Properties : Cell
		 Region : Cell 		Area
				Roundness
3. Population selection
 Population : Lyso 	Method : Filter by Property Output Population : Lyso Selected
	Cell Area [µm²] : > 5
	Cell Roundness : > 0.45
	Boolean Operations : F1 and F2
4. Define results
Method : List of Outputs
Population : Lyso Selected
	Number of Objects
	Apply to All :
	Intensity Cell Exp2Cam1 Mean :
	Intensity Cell Exp2Cam1 Sum :
	Cell Area [µm²] :
	Cell Roundness :
Population : Lyso Selected : ALL
Fig. S1B. Rosazza et al.
