## Supplementary figures and images for "Dynamic High-Content Imaging Reveals Surface Exposure of Virulent *Leishmania* Amastigotes in Infected Macrophages Undergoing Pyroptosis"

### Supplemental figure 2

#### Slide 1
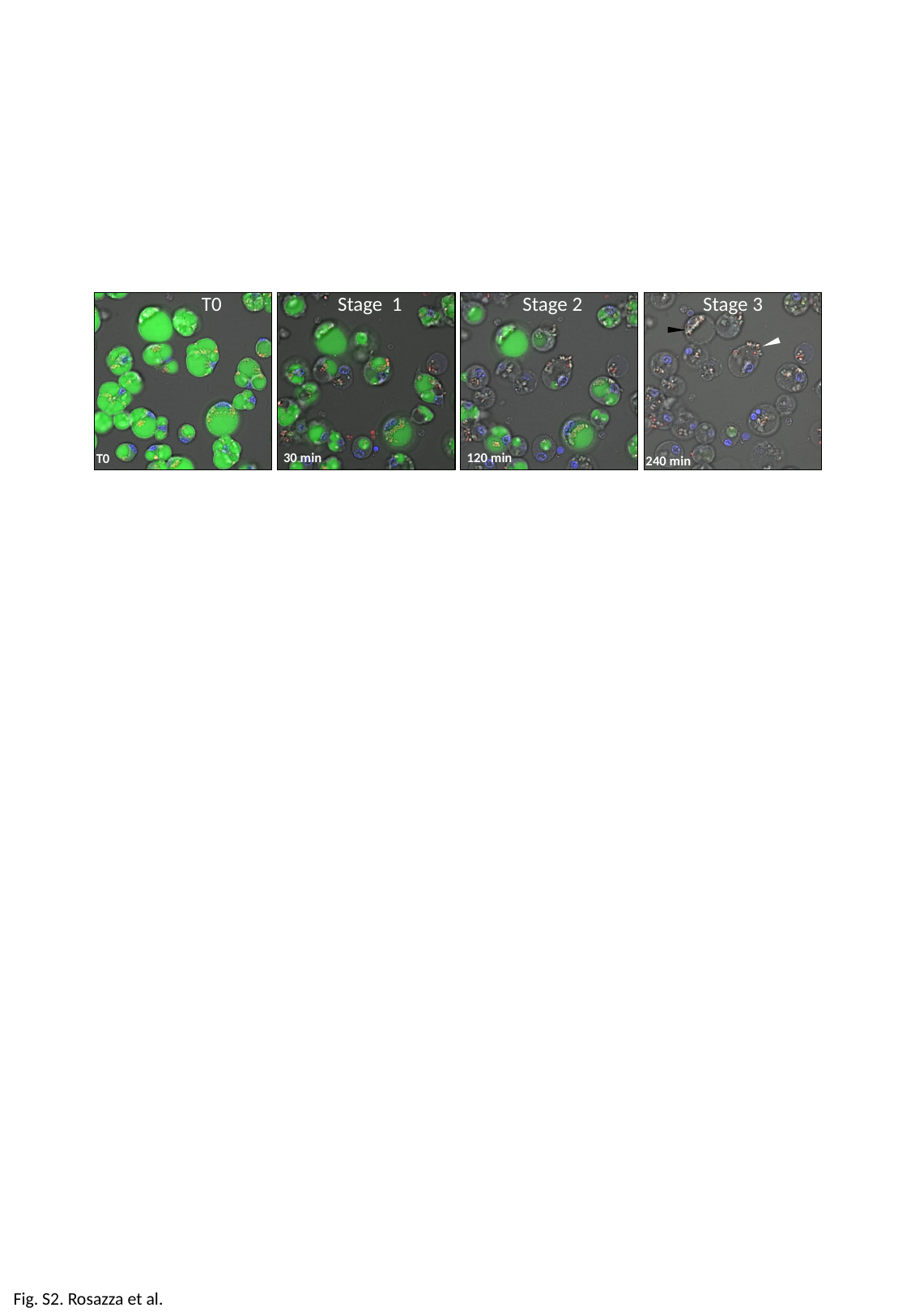

T0 Stage 1 Stage 2 Stage 3
30 min
120 min
T0
240 min
Fig. S2. Rosazza et al.

### Supplemental figure 3

#### Slide 1
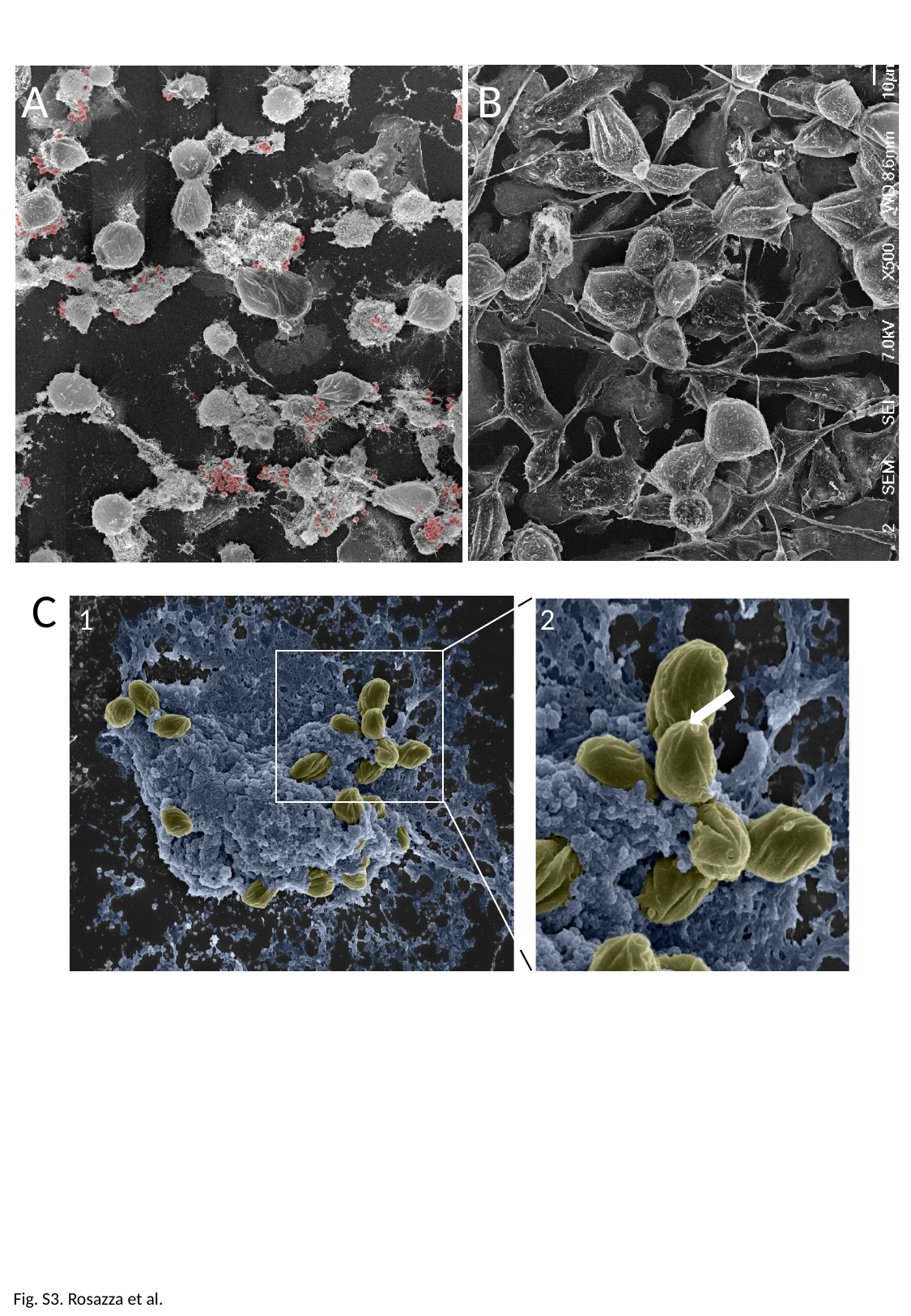

A B
 C
1 2
Fig. S3. Rosazza et al.
