## Supplemental figure 4 for "Dynamic High-Content Imaging Reveals Surface Exposure of Virulent *Leishmania* Amastigotes in Infected Macrophages Undergoing Pyroptosis"

### Slide 1
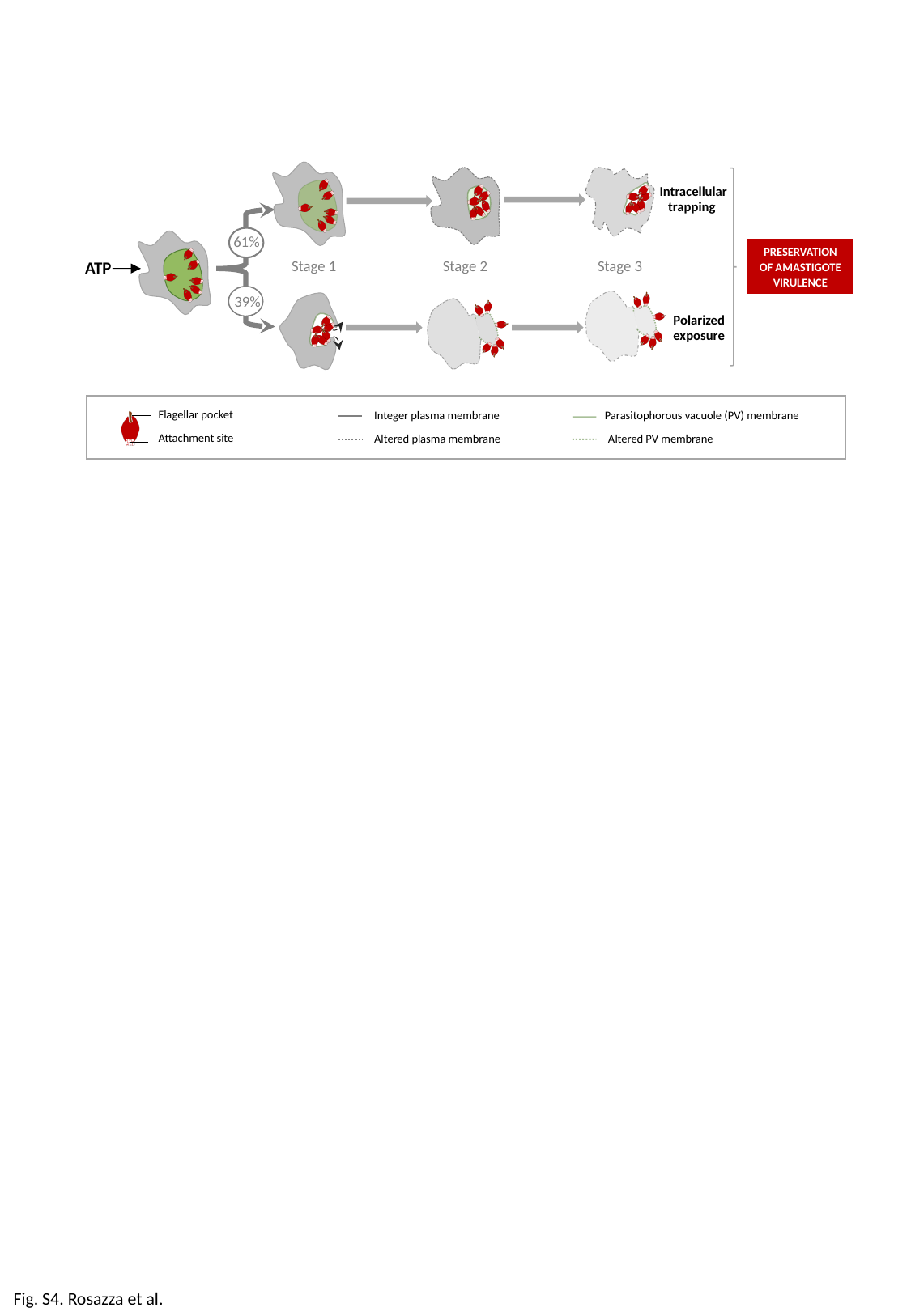

Intracellular trapping
61%
PRESERVATION OF AMASTIGOTE VIRULENCE
Stage 1 Stage 2 Stage 3
ATP
39%
Polarized exposure
Flagellar pocket
Attachment site
Integer plasma membrane Parasitophorous vacuole (PV) membrane
Altered plasma membrane Altered PV membrane
Fig. S4. Rosazza et al.
